# Set-up, validation, evaluation, and cost-benefit analysis of an AI-assisted assessment of responsible research practices in a sample of life science publications

**DOI:** 10.64898/2026.01.23.701317

**Authors:** Silke Kniffert, Ben Katthöfer, Robert Emprechtinger, Pasquale Pellegrini, Eva Maria Funk, Ishminder Singh Dhamrait, Yalei Zang, Ailyn Bornmüller, Ulf Toelch

## Abstract

The (semi-)automated screening of publications for diverse quality and transparency criteria is at the core of systematic literature assessment. Typically, the assessment process involves two initial reviewers and one additional reviewer for cases that require reconciliation. Here, we explore to what extent this process can be assisted by Large Language Models (LLMs). Specifically, whether LLMs are capable of assessing responsible research practices (RRPs) in scientific papers in a robust way. We employed proprietary LLMs to assess an initial set of 37 papers across ten RRPs. The same papers were also reviewed by three human reviewers. We iteratively redesigned prompts to increase model accuracy compared to human ratings which we treated as the gold standard. The resulting pipeline was validated on an additional set of 15 papers. We show that LLM accuracy is comparable to single human reviewer performance (90% for LLM vs 86% for a single human reviewer). However, performance strongly depended on the specific RRPs with accuracy ranging from 40% to 100%. LLMs exhibited an affirmative bias, making more errors when practices were not reported in the papers. Overall, we show how such an approach potentially replaces one human reviewer, enabling AI-assisted assessment of research papers. We discuss how dataset imbalances, validation procedures, and implementation time limit the broad applicability of such approaches. Through this, we develop initial guidance on the utility of proprietary LLMs in evidence synthesis.

## Introduction

Evidence synthesis constitutes an integral part of the cumulative scientific enterprise. The systematic identification, assessment, and integration of extant evidence serve to consolidate current knowledge and delineate research gaps. It thus informs practices and regulations in diverse fields, such as healthcare, business, and policy (Kunisch et al., 2023; Snyder, 2019). A central process in such work is the extraction of qualitative and quantitative measures from publications. As research reports in many fields are increasing in speed and volume, evidence synthesis becomes more critical, yet also time-consuming. Lengthy assessment and extraction processes delay the dissemination of important insights and, in fast-moving fields, potentially render a synthesis obsolete by the time of publication.

Potential support comes from the recent developments in the field of AI. Particularly, Large Language Models (LLMs) have already been tested and employed to support evidence synthesis. For example, the eligibility screenings for the selection of studies benefit from LLM support (Delgado-Chaves et al., 2025; Ghossein et al., 2025; Guo et al., 2024). Beyond title and abstract screening, full-text assessment and data extraction have been tested (Gartlehner et al., 2024; Li et al., 2025; Mitchell et al., 2025; Rahgozar et al., 2025; Woelfle et al., 2024) as well as trustworthiness assessment in clinical trial reports (Au et al., 2025). Moreover, AI potentially benefits more complex risk of bias assessments (Chao et al., 2024; Forero et al., 2025; Šuster et al., 2024; Xia et al., 2025) or adherence to reporting guidelines (Wrightson et al., 2024). Reviews of this emerging literature report an increase in studies utilising AI, primarily in a human-in-the-loop application scenario, as standalone AI remains error-prone (Clark et al., 2025).

As with many novel methodologies, there is currently no consensus on good practices to ensure that evidence generated with the assistance of AI is robust (Dijk et al., 2023). This is particularly important, as many LLMs available through cloud-based services are black-boxed. The black box dilemma poses a challenge for evaluating LLMs in assessment procedures, such as systematic reviews. One could use existing systematic reviews that humans have reviewed as a basis for estimating the accuracy of LLMs.

However, as existing systematic reviews have potentially already been used in the training of Large Language Models, they cannot serve as a basis for evaluating the capabilities of LLMs. That is, answers by LLMs potentially contain assessments by human reviewers, and high accuracy is not necessarily the product of reasoning processes of the LLM. There is currently no hard evidence on this as training datasets are usually not disclosed, but we deem it likely that the scientific literature is used extensively in AI training. Moreover, even if LLMs were trained on human assessments, it is not clear whether the LLM answers will recapitulate those. Therefore, at the onset of each project, LLM answers need to be validated on a novel dataset, a time-consuming process. As the field is relatively young, validation processes, including validation dataset size and techniques are not fully explored yet. Post-validation, implementation in a controlled screening environment is essential to assess the tool’s practical utility and real-world performance.

Here, we report on a project that extracts a set of Responsible Research Practices (RRPs) from a body of literature from the life sciences. RRPs foster trustworthiness of research procedures and results. Moreover, they contribute to the usefulness of research for researchers and society (Strech et al., 2020). Examples include strategies to minimise biases, such as randomisation and blinding, or transparent reporting, like open data and code. Assessment of reporting quality is predicated on RRPs, which are indispensable for ensuring the methodological validity and practical utility of the synthesized evidence. The criteria thus have a large overlap and extend criteria usually assessed in systematic reviews.

Here, we describe the process of implementing LLMs in the assessment process. We asked whether LLMs can increase the efficiency of assessment processes regarding RRPs in publications. We hypothesized that answers from LLMs show equal or superior accuracy compared to those of a single human reviewer. This would allow the partial replacement of human reviewers in such assessment processes and increase efficiency qua overall time and human review effort.

## Methods

To test our hypothesis, we first prioritized a set of RRPs for biomedical research through a Delphi consensus. We subsequently evaluated RRPs on a subset of life sciences papers stemming from our complete education assessment sample. Through systematic assessment by two human reviewers that included a reconciliation process, we created a novel dataset on the application of RRPs in the subsample. We utilized part of this dataset to iteratively design prompts that would increase the number of correct answers by different cloud-based commercial LLMs. This was streamlined through a pipeline for automated Application Programming Interface (API) calls to the respective LLM services. We validated our prompt in the remaining part of our human reviewed dataset. We retrospectively estimated costs for training and review processes to give guidance on which projects will benefit from such AI-assisted pipelines (Figure 1).

**Figure 1.**
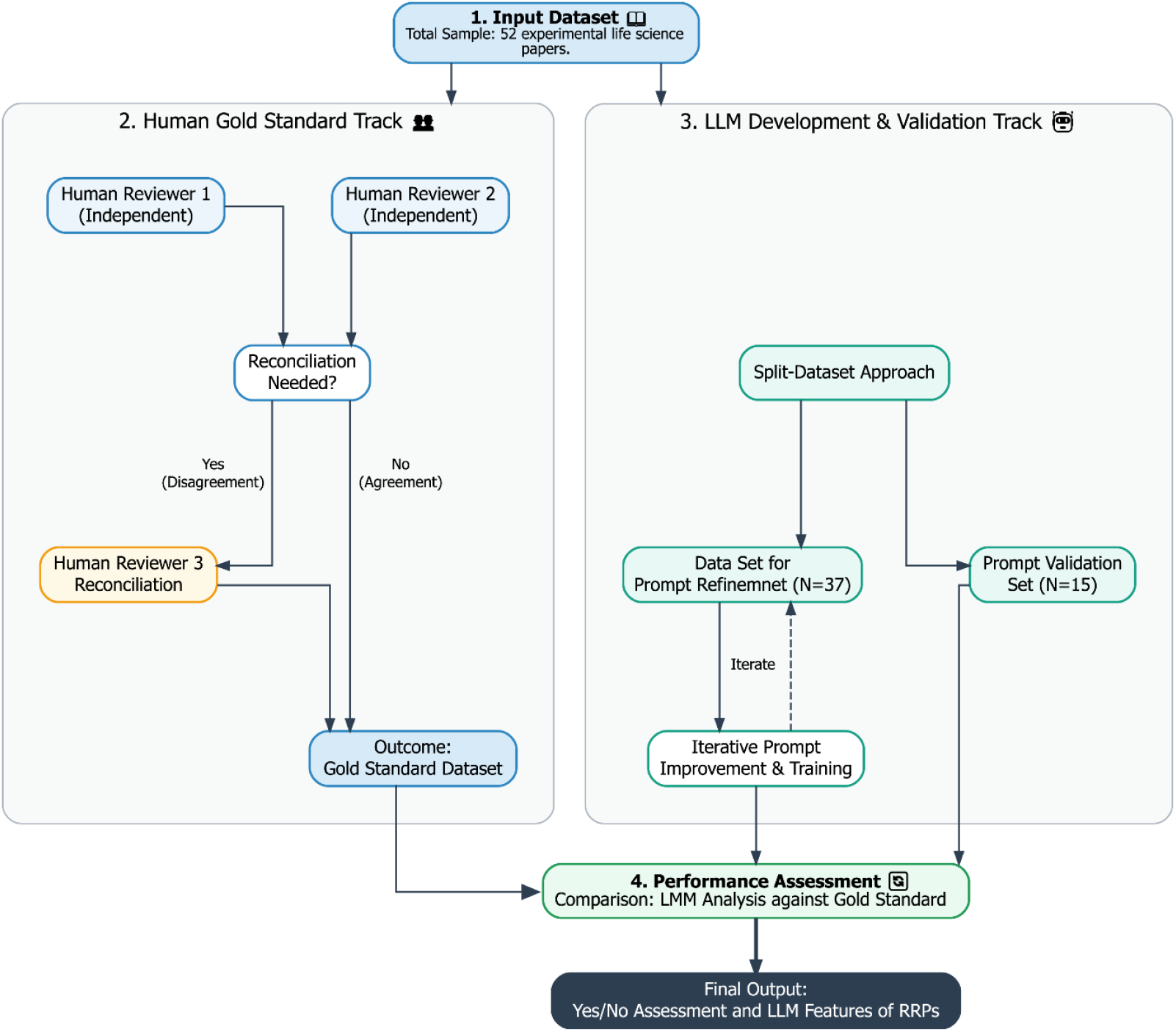
Assessment process of Responsible Research Practices by human reviewers and LLMs. A total of 52 experimental papers were assessed by two human reviewers. If case assessment differed between reviewers, a third reviewer did a reconciliation. This final assessment served as gold standard to assess accuracy of the initial two reviewers. LLM performance was tested on a dataset of 90% of the papers and prompts were improved until LLM performance reached approximately the accuracy level of a single reviewer. A hold out dataset of 10% was used to validate generalizability of the LLM assessment process.

### Delphi Study

In a first step, we aimed to prioritize a set of RRPs that would be most important for the quality of a paper. To operationalize the concept of RRPs, we first conducted an inventory of the learning objectives from the Berlin-Oxford Summer School on Responsible Research (BOX) (Gerrits et al., 2022; Toelch & Ostwald, 2018). BOX is a free to attend, international training event dedicated to guiding early-career researchers toward open, transparent, and reproducible research practices in the life sciences including psychology. It offers an introduction to responsible research workflows, helping attendees develop skills with a lasting impact on their careers and scientific communities. Based on this foundation, we developed a taxonomy categorizing RRPs into three overarching dimensions as a framework for RRPs: Inter- and Transdisciplinarity, Scientific Rigor, and Legal Rights. The dimension of Scientific Rigor was further divided into three subdimensions: Ethical Rigor, Methodological Rigor, and Sharing and Dissemination. In an iterative process with experts from the biomedical sciences and the monitoring and evaluation unit, teaching materials from the BOX were mapped to these thematic areas in several consensus meetings with experts, which enabled us to derive measurable indicators of responsible research. The thematic mapping process was primarily informed by structured expert elicitation, drawing upon collective professional judgment. In instances where initial consensus was not achieved, definitive categorization was subsequently guided by a targeted review of the extant literature and iterative deliberative refinement. This process resulted in a set of 49 Responsible Research indicators (Table S4).

In our Delphi study, the subdimension *Methodological Rigor* was selected as the primary focus, which reflected its central role in the thematic orientation of the BOX. For this, we defined criteria related to the knowledge base, validity and robustness, data management, documentation, translation, and sensu transfer of basic research findings into patient benefit. To identify the ten indicators most relevant to responsible scientific reporting, we conducted a three-round Delphi study. In the first round, a survey was distributed within our institute and further disseminated through snowball sampling.

Snowball sampling was employed by inviting participants to distribute the survey within their professional networks. Participants were asked to identify and rank the RRPs indicators they considered most important for responsible research reporting. In the second round, we held a consensus meeting with subject-matter experts to address conceptual ambiguities and incorporate feedback from the first round. Based on this discussion, the survey instrument was revised, including refined indicator definitions and updated question formats to facilitate faster responses and more robust analysis. The third round used the adapted survey and a revised dissemination strategy to finalize the selection of the top ten indicators.

### Selection of experimental research papers and human review

This project is part of a larger endeavour to estimate the impact of higher education courses on research outputs of participants and the application of RRPs. In this project, we collected publications of participants of the BOX. We also included in our sample publications from three control groups resulting in a sample of approximately 300 publications. For this study, we randomly selected 52 experimental papers (37 for prompt training and 15 for validation). We initially aimed for 40 papers in the prompt training, but three papers were excluded as they were systematic reviews. These numbers are to a certain degree arbitrary, but we aimed for an approximate 90%/10% training/test split. This deviates from the usual Pareto split in machine learning (ML) scenarios, combined with cross-validation through many different splits between training and test data. In our case, however, we aimed to refine the prompt to a fairly large part of our sample and then test for performance differences in a hold-out data set. The reasoning behind this is that we did not change the algorithm or weights of the LLM in the training but only the prompt. Consequently, we did not expect that our initial dataset would influence the outcome for subsequent datasets as severely as in ML training. We acknowledge this open issue of the need for training/test data for prompt evaluation in the discussion.

All papers (excluding supplements) were also reviewed by at least two human reviewers. Conflicts between human reviewers were resolved by a third reviewer to arrive at a consensus assessment. The human assessment was used as the gold standard that LLM results were compared to.

### Prompt Design

We created an initial prompt with an introductory part and then ten questions regarding RRPs (Table 1). The question about the tested population (Q1-3) was divided into three parts, one question for human, animal, or cell culture experiments. The question regarding the statistical rigour (Q7) was also divided into three sub-questions asking whether the test, test statistics and statistical decision criterion (like p-value) were reported. The latter question was later pooled into one answer. If any of the three questions indicated a “no” the study was rated as not appropriately reporting statistics. For prompt training, we started off with minimal questions (e.g. *Were experimental units or participants randomised to treatment groups?*). We then identified papers in the dataset with low accuracy and engaged in a prompt improvement process for the publications. We repeated the query and asked why the LLM answered in a certain way, asking for specific passages that elicited the response. We then asked the LLM how the prompt could be refined to improve answers on incorrect items. This process was repeated several times per publication until we could no longer improve the accuracy by this simple technique. This yielded often more complex and nuanced sub-questions that took also into account cases where this item was not applicable to the specific study (e.g. Q11): *Does the paper explicitly mention that animals or participants were randomly assigned or allocated to different experimental groups? If yes, report if and what method was used for randomisation. If no, mention if this was an observational study or a within subject design*.). This prompt improvement was done only on the Gemini model and for a limited number of iterations (max 10). With this, we do not know whether we reached saturation and if further improvement of prompts will lead to higher accuracy. The same prompt derived from Gemini was submitted to all models. See Supplemental Material for final prompt.

**Table 1.**
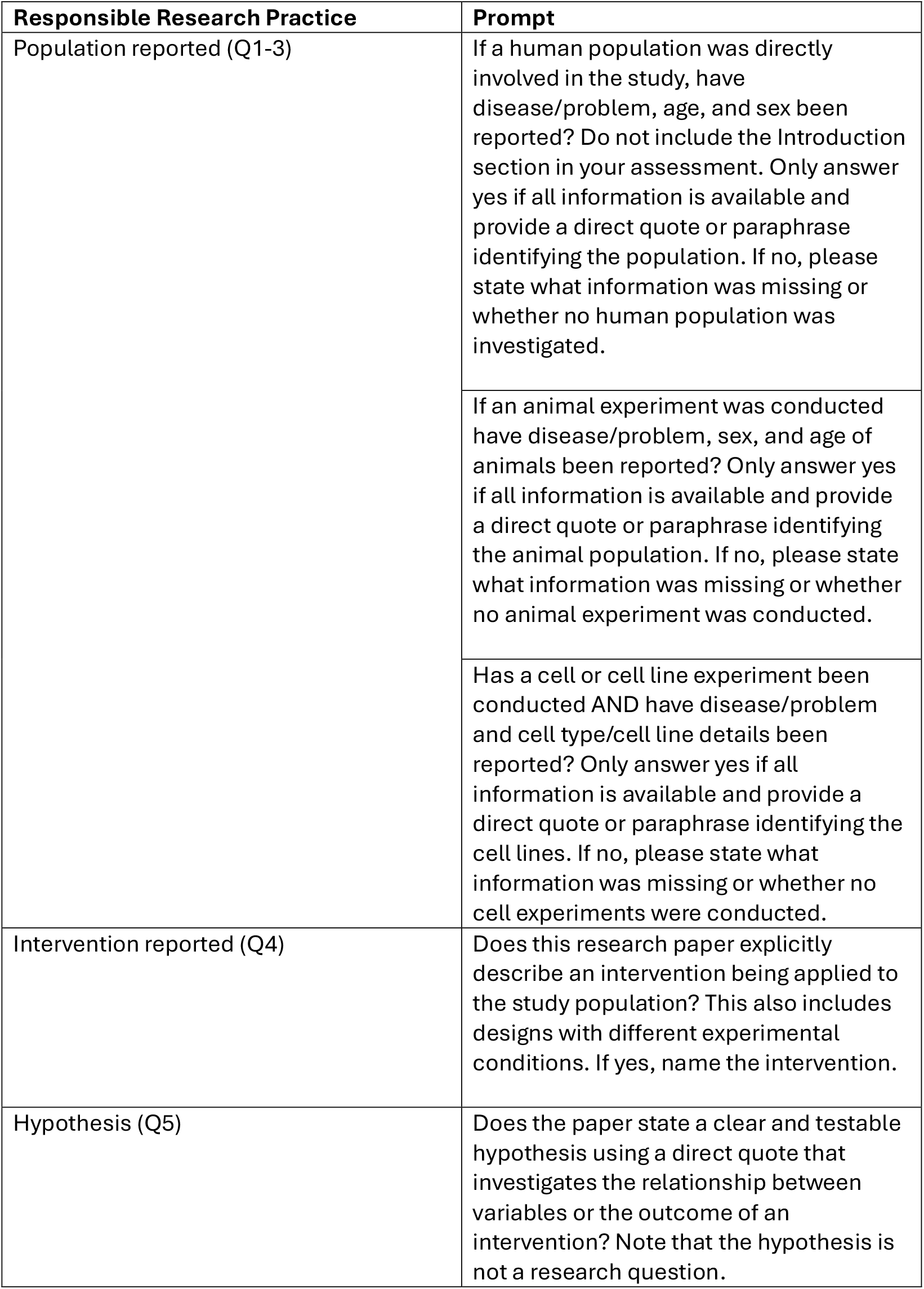
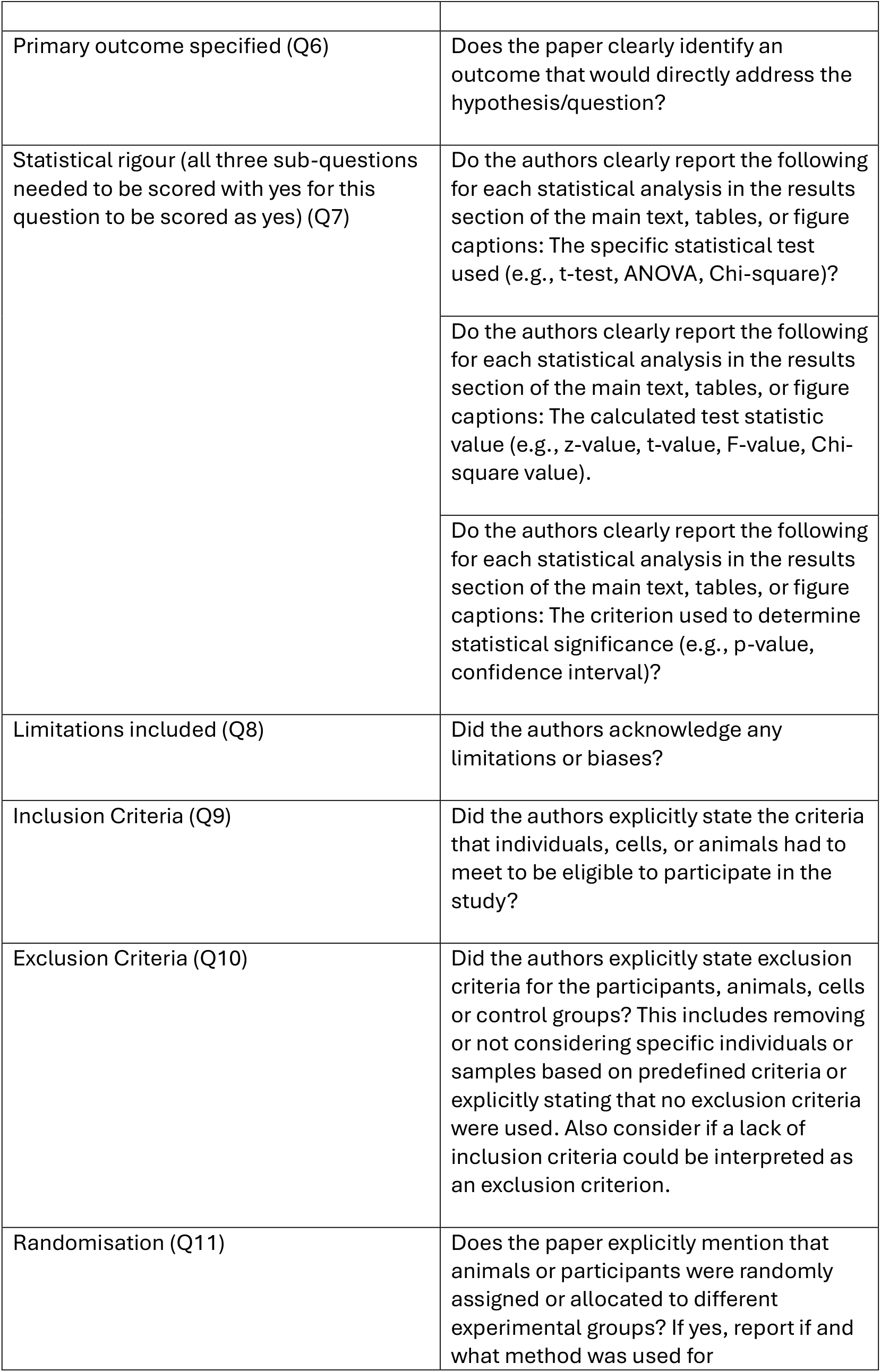
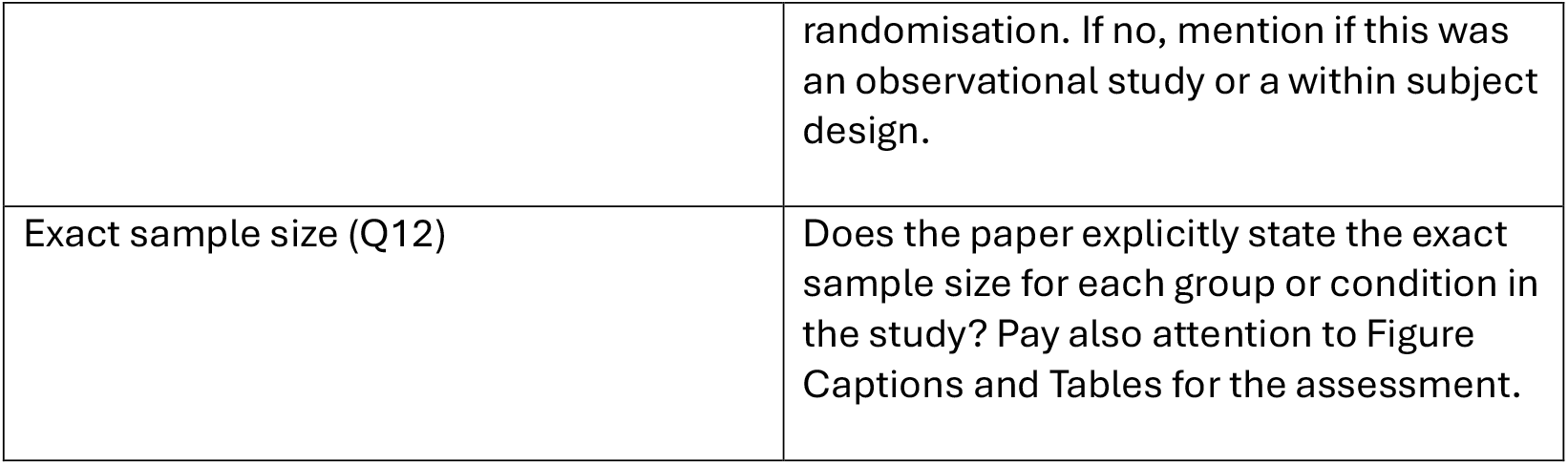
Responsible Research Practices assessed in the papers and the associated final prompt.

| Responsible Research Practice | Prompt |
| --- | --- |
| Population reported (Q1-3) | If a human population was directly involved in the study, have disease/problem, age, and sex been reported? Do not include the Introduction section in your assessment. Only answer yes if all information is available and provide a direct quote or paraphrase identifying the population. If no, please state what information was missing or whether no human population was investigated. |
|  | If an animal experiment was conducted have disease/problem, sex, and age of animals been reported? Only answer yes if all information is available and provide a direct quote or paraphrase identifying the animal population. If no, please state what information was missing or whether no animal experiment was conducted. |
|  | Has a cell or cell line experiment been conducted AND have disease/problem and cell type/cell line details been reported? Only answer yes if all information is available and provide a direct quote or paraphrase identifying the cell lines. If no, please state what information was missing or whether no cell experiments were conducted. |
| Intervention reported (Q4) | Does this research paper explicitly describe an intervention being applied to the study population? This also includes designs with different experimental conditions. If yes, name the intervention. |
| Hypothesis (Q5) | Does the paper state a clear and testable hypothesis using a direct quote that investigates the relationship between variables or the outcome of an intervention? Note that the hypothesis is not a research question. |
| Primary outcome specified (Q6) | Does the paper clearly identify an outcome that would directly address the hypothesis/question? |
| Statistical rigour (all three sub-questions needed to be scored with yes for this question to be scored as yes) (Q7) | Do the authors clearly report the following for each statistical analysis in the results section of the main text, tables, or figure captions: The specific statistical test used (e.g., t-test, ANOVA, Chi-square)? |
|  | Do the authors clearly report the following for each statistical analysis in the results section of the main text, tables, or figure captions: The calculated test statistic value (e.g., z-value, t-value, F-value, Chi-square value). |
|  | Do the authors clearly report the following for each statistical analysis in the results section of the main text, tables, or figure captions: The criterion used to determine statistical significance (e.g., p-value, confidence interval)? |
| Limitations included (Q8) | Did the authors acknowledge any limitations or biases? |
| Inclusion Criteria (Q9) | Did the authors explicitly state the criteria that individuals, cells, or animals had to meet to be eligible to participate in the study? |
| Exclusion Criteria (Q10) | Did the authors explicitly state exclusion criteria for the participants, animals, cells or control groups? This includes removing or not considering specific individuals or samples based on predefined criteria or explicitly stating that no exclusion criteria were used. Also consider if a lack of inclusion criteria could be interpreted as an exclusion criterion. |
| Randomisation (Q11) | Does the paper explicitly mention that animals or participants were randomly assigned or allocated to different experimental groups? If yes, report if and what method was used for |
|  | randomisation. If no, mention if this was an observational study or a within subject design. |
| Exact sample size (Q12) | Does the paper explicitly state the exact sample size for each group or condition in the study? Pay also attention to Figure Captions and Tables for the assessment. |

### Preparation of PDFs

PDFs of the studies were downloaded from respective publisher’s webpages and stored locally. For OpenAI’s gpt models and GEMINI, PDF documents were uploaded in advance to a vector database and retrieved from there for the queries. For models where the API did not support file uploads, the text was extracted using pypdf v.5.1.0 and appended to the prompt (o1-preview). In the extracted texts URLs beginning with “www” or “http(s)” were removed. PDFs were submitted through a subscription/API token payment model to prevent LLMs from using queries for further training.

### LLM Integration

The LLMs are integrated through the corresponding API (Application Programming Interface). A query was submitted for each study, in which the prompt with the questions and the PDF of the study is provided. The prompt also contains the instruction to structure the answer in the format of a csv table. The following LLMs were used: gpt-4o, gpt-4o-mini, o1-preview, and gemini-1.5-pro. Different temperatures were also tested for the GPT and Gemini models. The temperature is a parameter in LLMs that controls how deterministic an output is. Lower temperatures will yield more deterministic, higher temperatures more creative answers. The tested temperature parameters cover the range of possible values. A temperature parameter of 1.0 is the standard in most LLM agents. We tested more (<1.0) and less (>1.0) deterministic temperatures to cover a broad range of possibilities. We provide information on all runs and days when assessment was conducted (Table S2). In one case, we performed two runs with equal temperatures to see whether there were noticeable differences between models. LLM models were selected via an availability heuristic. We acknowledge that there are many models and parameter combinations available that could have been included. Our overall conclusions will thus be limited to these models and may not generalize to all models and parameter values.

### Study Evaluation Process

For evaluation, the raw text from each answer was processed line by line and interpreted as csv. Subsequently, all answers were compared against an existing dataset reviewed by three human reviewers (Figure 1). Answers included a statement whether the criterion was fulfilled and a feature that was extracted from the paper or was given by the LLM to justify the answer. The accuracy of a run is calculated as a percentage of correctly matching responses. Sensitivity/Recall (True Positives / (True Positives + False Negatives)), specificity (True Negatives / (True Negatives + False Positives)), precision (True Positives / (True Positives + False Positives)), and F1 values (2 * (Precision * Recall) / (Precision + Recall)) were calculated to assess model performance.

### Analysis

Data were pre-processed in Python (RRID:SCR_008394) code is provided at: https://doi.org/10.5281/zenodo.17209226. Data analysis was conducted in R (RRID:SCR_001905).

### Efficiency testing

If LLM performance is able to match the accuracy of human reviewers, we wanted to assess under what circumstances such a strategy can actually yield a time/resource benefit. For this, we recorded team members self-reported hourly investment into the assessment steps and the duration required for pipeline initialization and configuration. Through this we compared the time resources needed for validation process and LLM assisted assessment to human assessment alone. We extended this to different scenarios where assessment of a single paper needed either 20, 40, or 60 minutes of human review. For each scenario (minutes needed per paper) and number of papers (up to 500 papers in steps of 50), we simulated 1000 review processes. Time per paper was modelled as a Poisson process with the respective mean time as lambda.

## Results

### Results: Delphi

In the final round of the Delphi survey, 110 participants rated 49 methodological indicators. As not all respondents selected a full set of ten indicators (78.2% or n = 86/110), calculation of medians was not feasible. To account for this, ranking points were assigned to each indicator. Table S1 in the supplemental material presents the ten indicators with the highest ratings. Each indicator received between 20 and 32 ratings from participants. The *detailed research question* received the highest score (sum score= 254; n = 31), followed by the *hypothesis* (sum score= 182; n = 24). The remaining indicators in the top ten achieved mean scores between 143 and 114. The lowest score within this group was assigned to *sample size* (sum score= 114; n = 24).

### Results: Human and LLM assessment

Human reviewers achieved an individual accuracy of 87% that led to a reconciliation process in 25% of the cases. For LLMs, when submitting the revised prompt with twelve questions (10 RRPs) to four different LLMs, it resulted in above-chance accuracy in all models (>78%). The highest accuracies were obtained from Gemini when reducing the temperature to 0.2 followed by a reasoning model (o1-preview), both achieving accuracy values of ∼90% (Figure 2A). For cost reasons, we ran only Gemini and gpt-4o with different temperatures. To obtain these values, we had to refine the prompt over a maximum of ten consecutive runs (with minor adaptations in between). The initial prompt scored an accuracy of 81%. In the following, we focus on the highest accuracy model gemini-1.5-pro with a temperature of 0.2.

**Figure 2.**
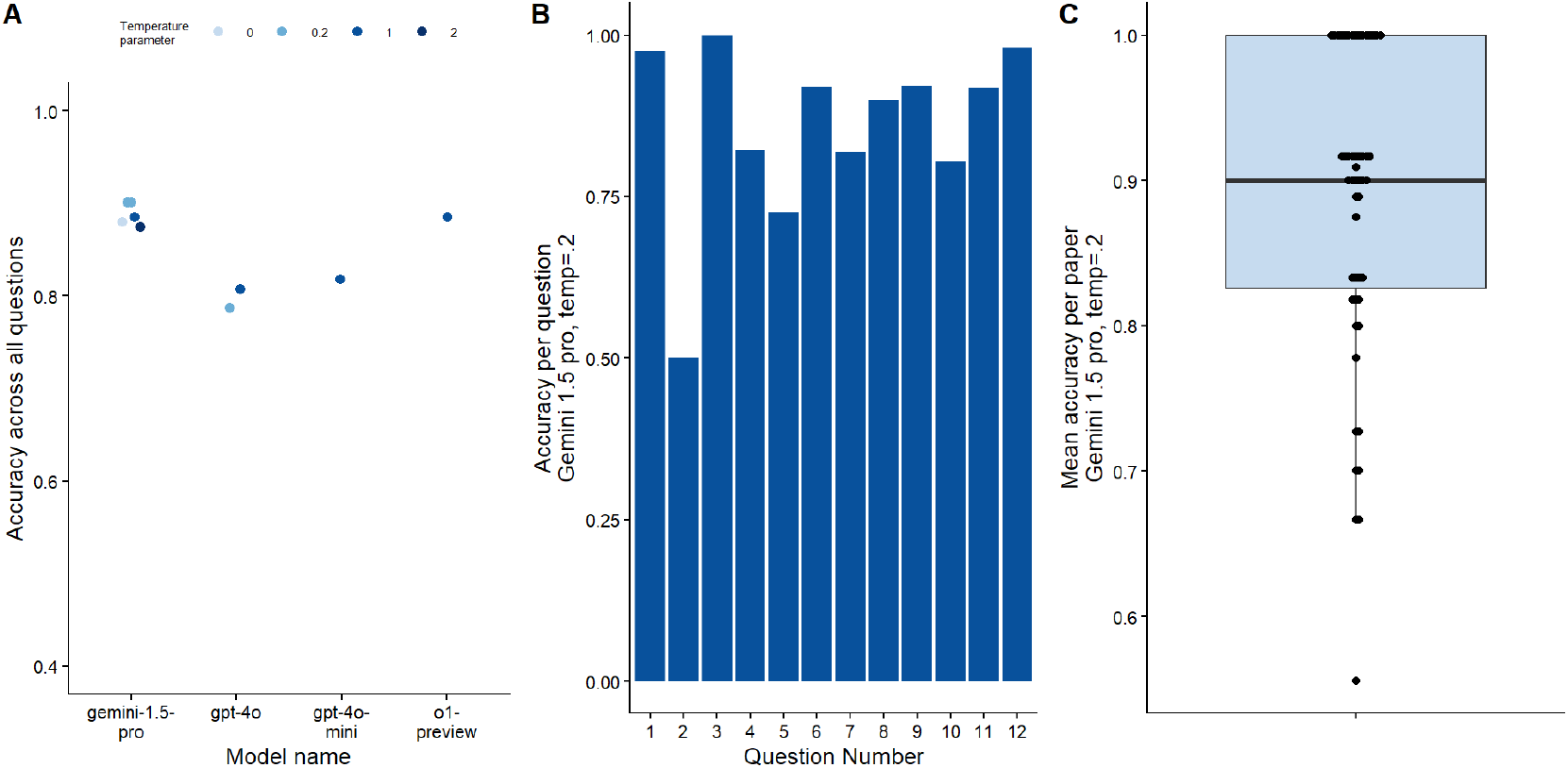
Comparison of accuracy. A. Accuracy for each tested model under different temperature parameters. Data includes only training dataset (n =37 papers) B. Accuracy for Gemini 1.5 model (Temperature Parameter = 0.2) for each of the different questions (n paper=52). For question numbers see Table 1 C. Accuracy of the Gemini model across papers. Each dot signifies one paper in the dataset (n paper=52).

Across questions there were marked differences in accuracy. For animal studies only 50% of questions regarding the population reporting were answered correctly (question 2). For other questions, like the proper reporting of the primary outcome (question 6), accuracy was higher than 90% (Figure 2B). Overall, the range was from 50 to 100% correct answers. Across papers, accuracy also differed with high accuracy scores in many papers above 90%, while others showed accuracies as low as 56% (Figure 2C).

Answers were not dependent on a specific iteration as a repeated submission of the same prompt yielded the same accuracy across all questions. Some questions did not apply as for example randomisation in observational studies. We excluded these questions based on the human reviewer assessment. To nonetheless assess the LLM performance in such cases its ability to rate a question as not applicable, we analysed the feature LLMs returned. These contained either a quote from the text or reasoning why the answer was negative. For all cases where a question did not apply, we rated (single reviewer) the statements and found that in 94% of the cases, the LLM gave a valid reason for the question to be excluded (see Table S3).

Next, we split papers into two answer groups, in which RRPs were either not detected (“no”) or detected (“yes”). Across questions, accuracy was lower in cases where the correct answer was “no” compared to “yes” answers (mean yes = 0.98; mean no = 0.67; two-sample test for proportions with continuity correction: χ^2^=93.6; df = 1; p < 0.0001; Figure 3A). This led to differential sensitivity, specificity, and precision across questions (Figure 3B). F1 values ranged from.57 to.99 and specificity from.5 to 1 (Figure 3B).

**Figure 3.**
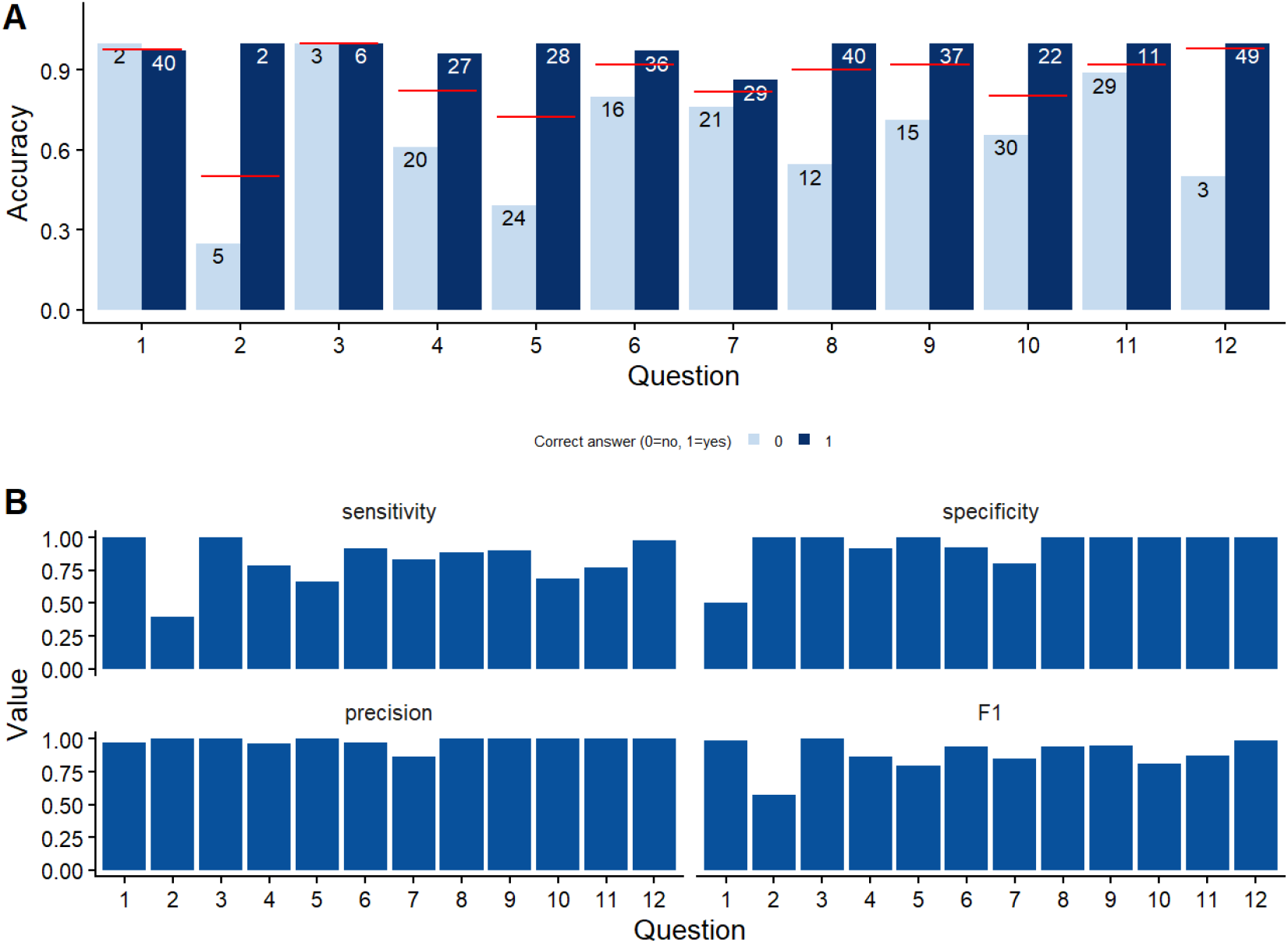
Evaluation metrics for Gemini 1.5 model (Temperature Parameter = 0.2). A. Accuracy for each question split by whether the paper implemented the RRP (‘yes’) or not (‘no’). Numbers indicate the number of papers in each category. Red line indicates the mean across yes/no answers. B. Sensitivity (recall), specificity, precision, and F1 for each question across all papers. Numbers per category are the same as in panel A. For question numbers see Table 1.

When compared with a single human reviewer from our team, most questions had similar accuracies (Figure 4A). Only for the RRPs regarding reporting of the animal population (Q2, Table 1), and whether a proper hypothesis was formulated (Q5 Table 1, Figure 4B) LLM performance was clearly lower (indicated by the two points above the line in Figure 4B). This resulted in a positive correlation between LLM performance and human accuracy for RRPs for each paper (Spearman rank correlation; rho=0.43; S=12621; p=0.0017). That is, if assessment was difficult for human reviewers LLMs were also less accurate.

**Figure 4.**
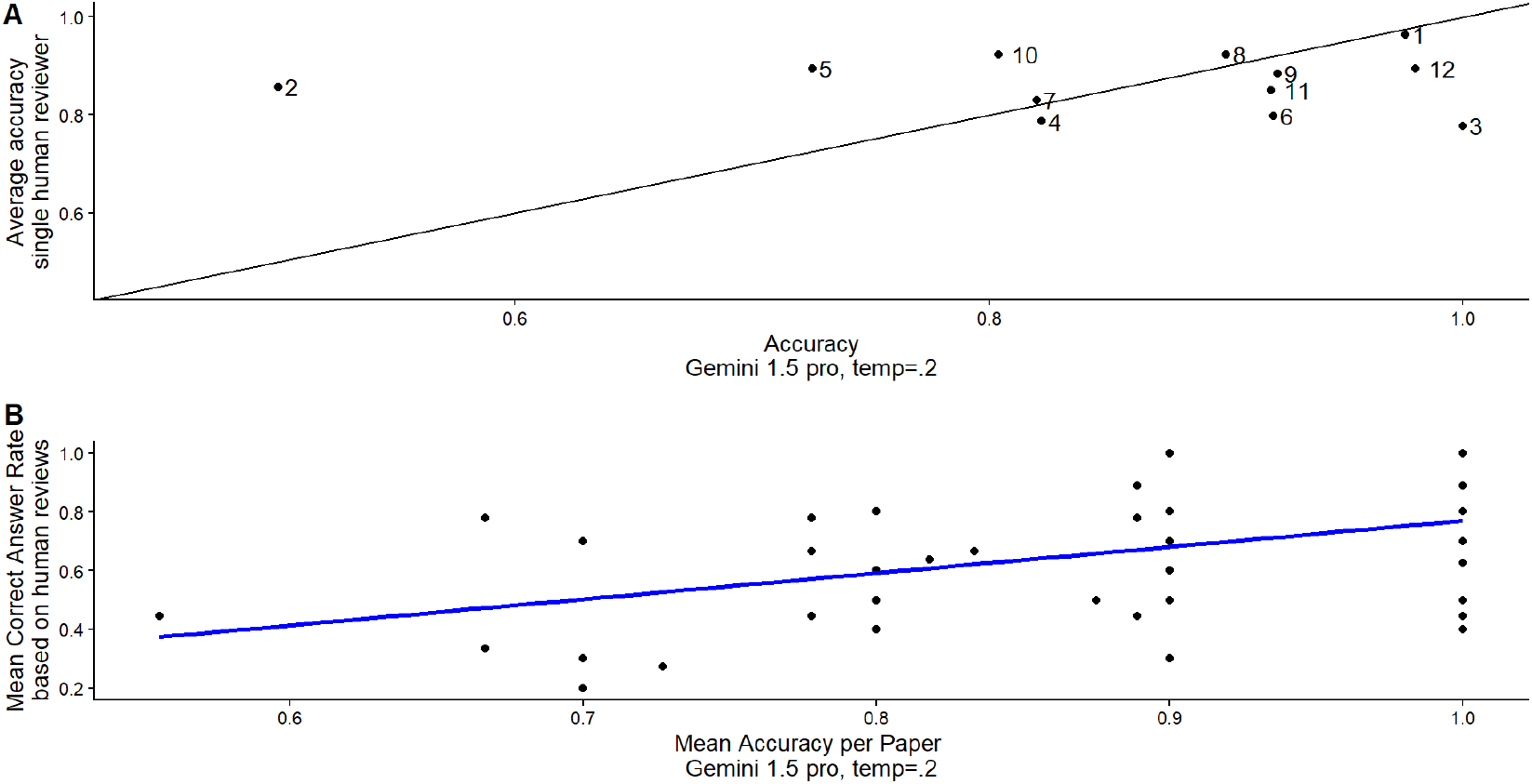
Correlation of accuracy of Gemini 1.5 model (Temperature Parameter = 0.2) with the average performance of a single human reviewer A. per question. and B. per paper.

### Results: Efficiency

We measured efficiency as total time needed to complete the review of a set of papers. The AI-assisted pipeline consisted of set-up, validation, and subsequent review. Setting up the pipeline to submit papers and prompt and receive feedback on the questions took approximately 50 hours. This does not include the time required to develop the pipeline itself, but only the time needed for set-up and initial tests. The number is based on an estimate by our data scientist (BK). The set-up included also time for refining the prompt and resubmission. Reviewers in our project needed 45 minutes on average for answering the questions for the selected papers (mean across six reviewers). For future tasks, we assume three scenarios, a slightly less demanding task where reviewers only need 20 minutes per paper and a demanding scenario requiring 1 hour per paper. We further assume for our simulation that reconciliation processes will be similar between AI-assisted and human reviewer only scenario. The net advantage of AI-based review processes is given by Eq.1. Note that time for training reviewers is not included here.

Equation 1: Advantage of AI-assisted review as a function of the number of papers (no_papers), average review time (avg_rev_time) and the time required for a set-up of the pipeline (set_up_time).

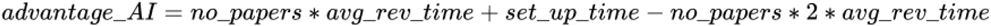

Our simulations show that, as expected, AI assistance is decreasing the total number of work hours for assessment if review times per paper are high or the number of papers to be reviewed is high (>100; Figure 5). If set-up times increase, the point of break-even will be even later. Cost-wise, we estimated that we spent 0.50, 1.53, and 15.63 Euro on a single run for the 37 initial papers for the Gemini 1.5, ChatGPT 4o, and o1 respectively (prices calculated in June 25). The efficiency gain of the AI is thus conditional on complexity of the task (time/paper) and scale (number of papers).

**Figure 5.**
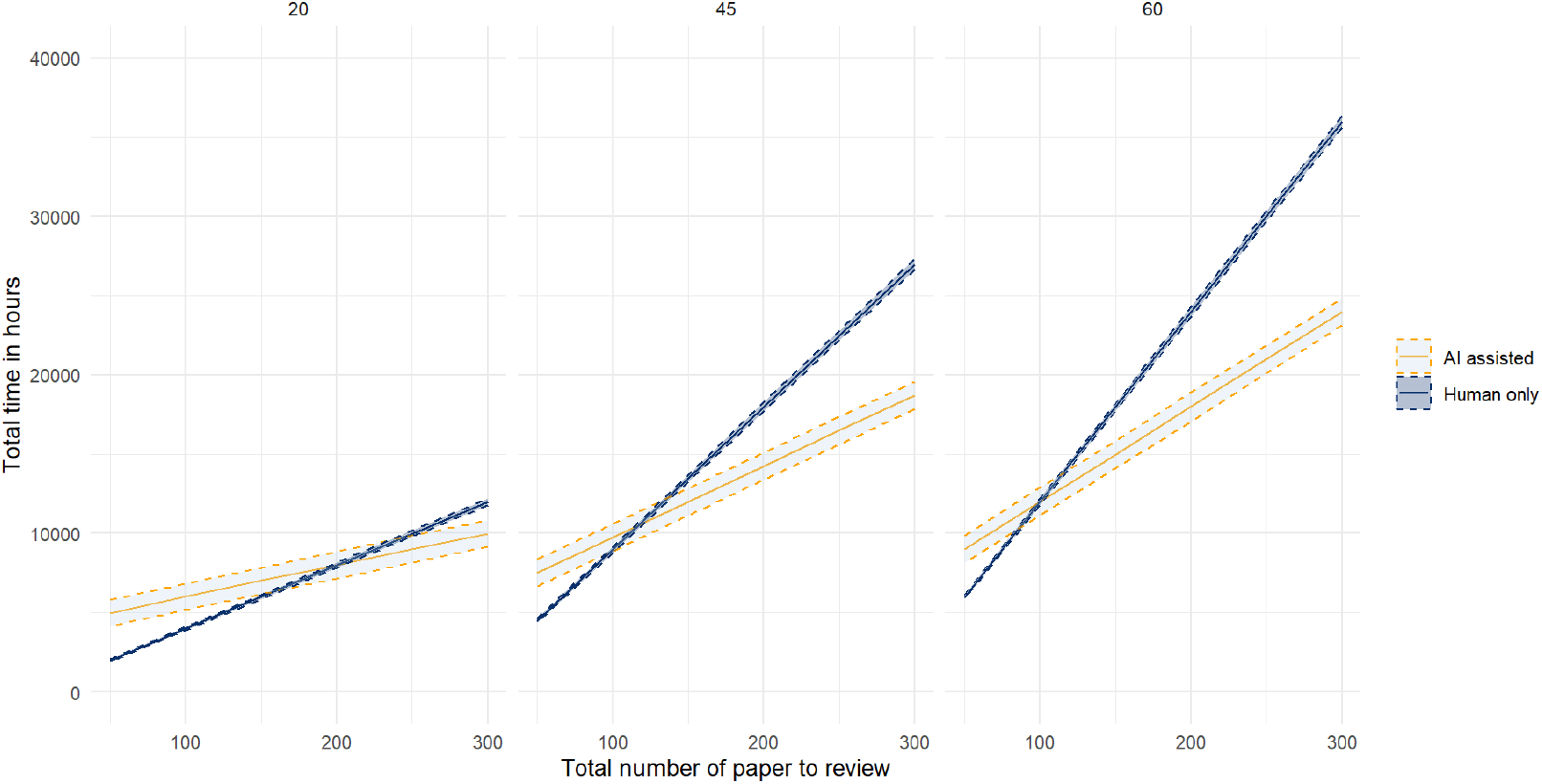
Comparison of time needed to review a set of papers in an AI-assisted scenario and in a human reviewer scenario. Individual panels refer to the time needed to review a single paper in minutes (20, 45, 60).

## Discussion

We probed whether use of general purpose LLMs meaningfully supports systematic assessment of Responsible Research Practices in a sample of papers from life science disciplines. We compared LLM performance on yes/no statements regarding RRPs to consensus of two human reviewers. After prompt optimisation, LLMs showed comparable performance to a single human reviewer. LLM answers, however, had an affirmative tendency showing lower accuracy when RRPs criteria were not fulfilled. This was especially true in a case in which questions only pertained to a small subcategory of paper or when the RRPs was especially complex to assess. Results of a utility analysis show that the decision to involve an LLM in such an assessment process depends on the scale of the project. For small-scale projects with only a small number of papers to be assessed, the effort for setting these processes up is potentially not warranted for economic and time reasons.

A range of limitations apply. These stem from the specific conduct of the Delphi study, the selection of models, and the validation and data quality.

The initial design of this study focused on the screening of publications within the biomedical sciences. Accordingly, the Delphi study to identify the most relevant RRPs indicators was conducted exclusively with biomedical experts. However, due to an insufficient number of eligible biomedical publications in our BOX sample, the scope of the screening was subsequently broadened to include studies from multiple disciplines. As a result, the RRP indicators derived from a discipline-specific (biomedical) Delphi process were applied to a cross-disciplinary corpus from the life sciences. This approach may have introduced a disciplinary bias, as the selected indicators may reflect priorities and norms more characteristic of biomedical research than of life sciences in general. Consequently, the generalizability of the selected indicators across disciplines may be limited.

We optimized the prompt on only one model (Gemini). This was partly motivated by the low (monetary) costs of this model, partly by the tier model of OpenAI where a minimum amount of money had to be spent to have access to additional models. Our results are thus biased against non-Gemini models and do not imply that similar accuracy levels could not be achieved for other model types. Particularly, we limited variation of the temperature parameter to Gemini and ChatGPT-4o. Moreover, prompt optimization was done manually. This resulted in a lack of reproducibility of the process by which one of the researchers (UT) arrived at the optimized version. Standardized, potentially automatized in interaction with LLMs processes could facilitate increased transparency and higher accuracy levels also for the optimization processes. During the development period, cloud-based models often changed sub-versions. This potentially impedes full computational reproducibility of results.

We optimized the prompt on overall accuracy across papers. In this process, we identified papers with low accuracy and adjusted prompts to increase single paper accuracy and then checked whether accuracy across all papers was increased.

Although this increased overall accuracy, the results are biased because the dataset was unbalanced. Some RRPs were more likely to be applied in a paper than others. That is, for some RRPs, the prompt was potentially biased towards “yes” answers. This is also reflected in analyses that calculate standard classification parameters like the F1 value and specificity. In the current version, the prompt did not yield high accuracy results in samples that do not apply RRPs. This potential issue arises not only from our unbalanced dataset but also from a general affirmative tendency in LLMs (Zhou et al., 2024).

Even if LLMs do not reach 100% accuracy, this is also a mark that human reviewers rarely achieve. Therefore, assessment is usually done by a minimum of two reviewers. We also employed such a human-in-the-loop scenario where LLMs are coupled with a human reviewer (Woelfle et al., 2024)It remains to be seen whether the combination of different LLMs or prompts will increase accuracy even further, but human involvement in such assessment processes remains a quality gold standard.

Beyond these limitations, LLMs can have a profound impact on the efficiency of systematic assessments, especially in large-scale projects such as systematic reviews, where implementation processes are time-consuming. Standardized, easy to implement pipelines may decrease the set-up cost. For example, in standard assessments like the PICOS scheme, LLMs increase efficiency in the assessment process (Vallamchetla et al., 2025). In our more specialized project, the number of papers that need to be reviewed has an influence on whether engaging with validation and prompt design is worthwhile. For our main project, that is not reported here, we aim to review over 300 papers. A number, projections show, that warrants additional implementation time for LLMs.

As assessment of, for example, the PICO scheme, risk of bias, etc., usually follows straightforward definitions, standardized large datasets and associated validated prompts that are created from the community for the community, potentially reducing implementation time (Bolaños et al., 2024; Kousha & Thelwall, 2024; Ofori-Boateng et al., 2024). The current disparity of approaches, our own included, precludes a common approach for AI-assisted meta-research. A standard toolbox with validated and standardized procedures would lower the threshold for users to apply such tools(Cao et al., 2025). With a focus on research questions rather than implementation assessment, speed will increase. This will help evidence synthesis to keep pace with the growing body of primary literature. As with any new technology, there is a need to clearly identify contexts where the application of LLMs will responsibly create utility.

## Acknowledgements

We thank all participants of the Delphi study and particularly participants in the consensus meeting: Natascha I. Drude, Maria Arroyo Araujo, Clarissa F.D. Carneiro, Charlotte Klein, and Maren Hülsemann

## Notes

### Competing Interest Statement

The authors have declared no competing interest.

### Funding

- Bundesministerium für Forschung, Technologie und Raumfahrt: 01KC1901A, 01KC2306
- Einstein Stitung Berlin: EZ&#8208;2020&#8208;597&#8208;2

