## Supplementary Material for "Set-up, validation, evaluation, and cost-benefit analysis of an AI-assisted assessment of responsible research practices in a sample of life science publications"

Supplemental material

Table S1 The ten indicators with the highest rating represented by the mean scores and the sum scores.

| | Indicators | Sum scores | Mean scores | N ratings |
|---|---|---|---|---|
| 1. | Detailed research question | 254 | 8,19 | 31 |
| 2. | Study protocol | 200 | 6,66 | 30 |
| 3. | Detailed method plan | 189 | 6,10 | 31 |
| 4. | Hypothesis | 182 | 7,58 | 24 |
| 5. | Analysis plan | 143 | 5,72 | 25 |
| 6. | Limitations | 141 | 4,41 | 32 |
| 7. | In- and exclusion criteria | 139 | 5,15 | 27 |
| 8. | Positive or negative control | 138 | 5,75 | 24 |
| 9. | Randomization | 122 | 5,08 | 24 |
| 10. | Sample size | 114 | 5,7 | 24 |

Table S2 Overview of different runs and parameters for the different models. The column validation run signifies whether the run was on the initial data set of 37 paper (False) or whether the run was on the hold out data set.

| ID | Model | Temperature | PDF-Reader | Validation-Run | Date |
|---|---|---|---|---|---|
| **1** | gemini-1.5-pro | 2.0 | - | False | 2025-06-08 |
| **2** | gemini-1.5-pro | 0.0 | - | False | 2025-06-08 |
| **3** | gemini-1.5-pro | 0.2 | - | False | 2025-02-26 |
| **4** | gemini-1.5-pro | 0.2 | - | False | 2025-04-25 |
| **5** | gemini-1.5-pro | 1.0 | - | False | 2024-10-06 |
| **6** | gemini-2.0-flash-exp | 0.2 | - | False | 2025-06-08 |
| **7** | gpt-4o | 0.2 | - | False | 2025-06-08 |
| **8** | gpt-4o | 1.0 | - | False | 2025-06-08 |
| **9** | gpt-4o-mini | 1.0 | - | False | 2024-10-08 |
| **10** | o1-preview | 1.0 | pypdf 5.1.0 | False | 2025-06-08 |
| **11** | gpt-4o | 1.0 | pypdf 5.1.0 | False | 2025-06-08 |
| **12** | gpt-4o-2024-08-06 | 0.2 | - | False | 2025-06-08 |
| **13** | gpt-4o-2024-08-06 | 1.0 | - | False | 2025-06-08 |
| **14** | gemini-1.5-pro | 1.0 | - | True | 2025-04-10 |
| **15** | gpt-4o-2024-08-06 | 0.2 | - | True | 2025-05-08 |
| **16** | gpt-4o-2024-08-06 | 1.0 | - | True | 2025-05-08 |
| **17** | gemini-1.5-pro-002 | 0.2 | - | True | 2025-06-06 |
| **18** | gemini-1.5-pro-002 | 1.0 | - | True | 2025-06-06 |
| **19** | gemini-1.5-pro-002 | 2.0 | - | True | 2025-06-06 |

Table S3 Features extracted from LLMs for assessing eligibility of question for the paper:

| pdf | Question number | explanation |
|---|---|---|
| 5 | 1 | The paper investigates the effect of Teriflunomide on mitochondria in the context of oxidative stress. The introduction mentions Multiple Sclerosis, but the study itself does not involve a human population. |
| 5 | 3 | The paper investigates the impact of Teriflunomide on mitochondria using explanted murine spinal roots. While this involves cells, it's in the context of a tissue model, not isolated cell lines. |
| 13 | 2 | No animal experiment was conducted. |
| 13 | 3 | No cell experiment was conducted. |
| 13 | 11 | This was an observational study. |
| 19 | 2 | No animal experiment was conducted. |
| 19 | 3 | No cell experiment was conducted. |
| 31 | 2 | No animal experiment was conducted. |
| 31 | 3 | No cell experiment was conducted. |
| 54 | 2 | No animal experiment was conducted. |
| 54 | 3 | No cell experiment was conducted. |
| 94 | 2 | No animal experiment was conducted. |
| 94 | 3 | No cell experiments were conducted. |
| 94 | 4 | This was an observational study. |
| 98 | 2 | The study did not involve any animal experiments. |
| 98 | 3 | The study did not involve any cell or cell line experiments. |
| 100 | 2 | No animal experiment was conducted. |
| 100 | 3 | No cell experiment was conducted. |
| 110 | 1 | No human population was investigated. |
| 124 | 2 | No animal experiment was conducted. |

| pdf | Question number | explanation |
|---|---|---|
| 124 | 3 | No cell experiment was conducted. |
| 129 | 2 | No animal experiment was conducted. |
| 129 | 3 | No cell or cell line experiment was conducted. |
| 172 | 1 | No human population was investigated. |
| 172 | 3 | No cell experiments were conducted. |
| 191 | 2 | No animal experiment was conducted. |
| 191 | 3 | No cell experiment was conducted. |
| 214 | 1 | No human population was investigated. |
| 214 | 2 | No animal experiment was conducted. |
| 223 | 2 | No animal experiment was conducted. |
| 223 | 3 | No cell experiment was conducted. |
| 223 | 4 | The paper does not describe an intervention. It investigates neuroanatomical markers. |
| 223 | 11 | The paper describes a longitudinal observational study design. |
| 226 | 2 | No animal experiment was conducted. |
| 226 | 3 | No cell experiment was conducted. |
| 280 | 2 | No animal experiment was conducted. |
| 280 | 3 | RNA was hybridized to Illumina HT-12 v4 Expression BeadChips (Illumina) and further processed as described in the Supplementary Materials. |
| 379 | 2 | No animal experiment was conducted. |
| 379 | 3 | No cell experiment was conducted. |
| 424 | 2 | No animal experiment was conducted |
| 424 | 3 | No cell experiment was conducted |
| 435 | 2 | No animal experiments were conducted. |

| pdf | Question number | explanation |
|---|---|---|
| 435 | 3 | No cell experiments were conducted. |
| 435 | 4 | The paper describes a qualitative study based on semi-structured interviews. |
| 435 | 11 | This was an observational study with a semi-structured interview design. |
| 480 | 2 | No animal experiment was conducted. |
| 480 | 3 | No cell experiment was conducted. |
| 491 | 2 | No animal experiment was conducted. |
| 491 | 3 | No cell experiment was conducted. |
| 535 | 2 | No animal experiment was conducted. |
| 535 | 3 | No cell experiment was conducted. |
| 541 | 2 | No animal experiment was conducted. |
| 541 | 3 | No cell experiments were conducted. |
| 541 | 11 | This was an observational study. |
| 646 | 1 | The study investigates the role of tau protein in anxiety and memory in mice. |
| 646 | 3 | No cell or cell line experiments were conducted. |
| 665 | 1 | The study did not involve a human population. |
| 665 | 2 | The study did not involve an animal experiment. |
| 705 | 2 | No animal experiment was conducted. |
| 705 | 3 | No cell experiment was conducted. |
| 714 | 2 | No animal experiment was conducted. |
| 714 | 3 | No cell experiment was conducted. |
| 732 | 2 | No animal experiment was conducted. |
| 732 | 3 | No cell experiment was conducted. |
| 732 | 4 | Participants experienced a single trial in one of the three driving conditions. In the first condition, a fully autonomous car with an anthropomorphic voice assistant system (AVAS) |

| pdf | Question number | explanation |
|---|---|---|
| | | provided information about critical traffic events and the corresponding car decisions. The second condition was an AV with a radio broadcast playing throughout the trial. In the third condition, a female TaxiDriver drove the participant through the city. |
| 732 | 11 | Participants experienced a single trial in one of the three driving conditions. |
| 760 | 2 | No animal experiment was conducted. |
| 760 | 3 | No cell experiment was conducted. |
| 819 | 2 | No animal experiment was conducted. |
| 819 | 3 | No cell experiment was conducted. |
| 819 | 4 | This was an observational study. |
| 819 | 11 | This was an observational study. |
| 837 | 2 | No animal experiment was conducted. |
| 837 | 3 | No cell experiment was conducted. |
| 887 | 2 | No animal experiment was conducted. |
| 887 | 3 | No cell experiment was conducted. |
| 891 | 2 | No animal experiment was conducted. |
| 891 | 3 | No cell experiment was conducted. |
| 935 | 2 | No animal experiment was conducted. |
| 935 | 3 | No cell experiment was conducted. |
| 94 | 7 | The authors do not report the specific statistical tests used. |
| 435 | 7 | No statistical tests were used, as this is a qualitative study. |

Table S4 Full List of indicators for the Delphi Study

| | **Indicators** |
|---|---|
| 1. | Detailed research question |
| 2. | Study protocol |
| 3. | Detailed method plan |

| | |
|---|---|
| 4. | Hypothesis |
| 5. | Analysis plan |
| 6. | Limitations |
| 7. | In- and exclusion criteria |
| 8. | Positive or negative control |
| 9. | Randomization |
| 10. | Sample size |
| 11. | Systematic literature search |
| 12. | Data repository |
| 13. | Preregistration |
| 14. | Standard operating procedure (SOP) |
| 15. | Blinding |
| 16. | Analysis pipeline |
| 17. | Sample size calculation |
| 18. | Primary outcome |
| 19. | Use of reporting guidelines |
| 20. | Statistical tests |
| 21. | Sex |
| 22. | Cell line |
| 23. | Research data management plan |
| 24. | Determination of the experimental unit |
| 25. | Age |
| 26. | Outlier handling |
| 27. | Disease mechanism |
| 28. | Source of funding |
| 29. | Statistical significance |
| 30. | Comorbidity |
| 31. | Precision |
| 32. | Effect size |
| 33. | Species |
| 34. | Attrition |
| 35. | Treatment dose |
| 36. | Use of an electronic lab notebook |
| 37. | Groups comparison |
| 38. | Treatment order |
| 39. | Baseline |
| 40. | Measuring time points |
| 41. | Power calculation |
| 42. | Day/ night cycle |
| 43. | Weight |
| 44. | Further environmental factors in lab or animal facility |
| 45. | Temperature in the animal facility |
| 46. | Genotype |
| 47. | Housing |

| | |
|---|---|
| 48. | Use of version control |
| 49. | Strain |
